# Active Inference as a Framework for Brain-Computer Interfaces

**DOI:** 10.1101/2021.02.02.429272

**Authors:** Syed Hussain Ather

## Abstract

As Karl Friston explained during the International Symposium on Artificial Intelligence and Brain Science 2020, active inference provides a way of using abstract rule-learning and approximate Bayesian inference to show how minimizing (expected) free energy leads to active sampling of novel contingencies. Friston elaborated how there were ways of making an optimal decision using active inference that can offer perspectives to advances in artificial intelligence. These methods of optimization within the context of active inference can also be used as a framework for improving brain-computer interfaces (BCI). This way, BCIs can give rise to artificial curiosity in the way Friston had described during his session. Using Friston’s free energy principle, we can optimize the criterion a BCI uses to infer the intentions of the user from EEG observations. Under Friston’s criteria for making an optimal decision, BCIs can expand their framework of optimal decision-making using active inference.

## 1. Introduction

### 1.1. Free energy principle

In the search for grand unified theories of the brain, the free energy principle states that a self-organizing principle at equilibrium with its environment must minimize the amount of free energy that it has. Its applications have spanned far and wide in explaining brain structure and function. This self-organizing behavior arises from the defining characteristic of biological systems to resist disorder in dynamic environments[1] stemming from Helmholtz’s ideas of perception[2]. A model of perceptual influence and learning built upon these ideas can explain how principles can resolve problems of the inference of the causes underlying sensory input and learning the causal structure that generates them. From there, one can study how inference and learning follow. Friston has used this in the context of Empirical Bayes and hierarchical models of sensory input to show how the free energy principle can understand a range of cortical organization and responses.

The free energy principle provides a way for adaptive systems to unify action, perception, and learning[3]. The theory and background of the principle involves a system using a Markov blanket to minimize the difference between a model of the world and its sense and the perception associated with it. Through continuously correcting the world model of the system, the system changes the world into an expected state while minimizing the free energy of the system. Using the Bayesian idea of the brain as an “inference engine,” the system can actively change the world (active inference) into the expected state and minimize the free energy of the system. This holds true for a wide variety of adaptive systems from animals to brains themselves in understanding mental disorders and artificial intelligence, and other applications of the free energy principle span areas of exploration and novelty seeking[4].

Friston outlined the motivation behind using the free energy principle as a unified brain theory using the system’s tendency to resist disorder. When a system resists its tendency to move towards disorder, the physiological and sensory states of a system move towards configurations of low entropy. Given that the number of these states is limited, the system is very likely to be in these states of low entropy. By using a formulation of entropy as the average amount of self-information or “surprise” (the negative log-probability of a specific outcome), Friston explained how biological agents minimize the long-term average of surprise (or maximize sensory evidence for an agent’s existence) to keep sensory entropy low. By sampling the environment to change configuration and minimize free energy this way, the system changes its expectations[5]. This forms the basis of action and perception, and the system’s state and structure encode an implicit and probabilistic model of the environment. The nervous system in particular maintains order through these methods, and the specific structural and functional organization is maintained by the environment’s causal structure.

One can evaluate free energy as a function of two things to which the agent has access: the sensory states and a recognition density encoded by its internal states (such as neuronal activity and connection strengths)[6]. In terms of the environment’s causal structure, the recognition density would be a probabilistic representation of what caused a particular sensation. The causes underlying sensory input, or the probabilistic representation of what causes sensations, are used as the recognition density. They can vary from an object in one’s field of vision or blood pressure changing the physiological state of organs.

### 1.2. Active Inference

A corollary of the free energy principle, active inference, arises from the way natural agents act in the context of these observations. Active inference claims natural agents act to fulfill prior beliefs about preferred observations. By adjusting sensory data (without changing recognition density) to minimize free energy, an agent chooses the sensory inputs from a sample based on prior expectations to increase the accuracy, the surprise about sensations expected under a particular recognition density, of an agent’s predictions. In the context of Bayesian inference, one may define the complexity (“Bayesian surprise”) as the difference between the prior density, which encodes beliefs about the state of the world before sensory data are assimilated, and posterior beliefs, encoded by the recognition density. In essence, the agent avoids surprising states by making active inferences.

As Friston explained during his session[7], active inference is self-evidencing in that action and perception can be cast as maximising Bayesian model evidence for the generative models of the world[8]. Using a generative model of the underlying causal structure of a system, you can explain how evidence accumulates or how a specific action was chosen. Examples of active inference for Markov decision processes include using Bayes optimal precision to predict activity in dopaminergic areas [9] and using a gradient descent on variational free energy to simulate neuronal processing [10]. This evidence can be decomposed through its accuracy and complexity.

One may even describe active inference as a method of explaining action using the idea that the brain has “stubborn predictions” that are resistant to change, such as adaptive body temperature necessary for survival, that cause the system to behave in a way to cause the predictions to come true[11]. Figuring out the etiology of stubbornness would provide insight into ways of how to change these predictions helpful for understanding the relationship between drugs and psychotherapy that have synergistic effects when used together. Other applications of active inference extend to visual foraging[12] and BCIs[13].

### 1.3. Active Inference Python Implementation

A good example of a Python implementation of Active Inference can be found in the “Active Inference for Markov Decision Processes” repository [14].

Using the `grid world` environment that has 3 x 3 states, an agent can choose one of five actions for each time step. The agent can move in any of the four directions (up, down, left, right) or remain where it is. After performing inferences about its current location, the agent must figure out which location it prefers to move to and move there. Using a reward value, we can set the reward value to reward the user when choosing the direction leading to a preferred location.

~~~
REWARD_LOCATION = 7
env = GridWorldEnv()
env.set_reward_state(REWARD_LOCATION)
~~~

In it, the “Inference and action example” illustrates the process of creating a generative model consisting of a likelihood distribution A and an empirical prior distribution B for the environment.

~~~
likelihood_matrix = env.get_likelihood_matrix()
A = Categorical(values=likelihood_matrix)
A.remove_zeros()
plot_likelihood(A)
transition_matrix = env.get_transition_matrix()
B = Categorical(values=transition_matrix)
B.remove_zeros()
plot_empirical_prior(B)
~~~

With the beliefs about the current location:

~~~
Qs = Categorical(dims=[env.n_states])
~~~

The beliefs are then plotted from the agent’s preferences:

~~~
C[REWARD_LOCATION] = 1.
plot_beliefs(C, title=“Prior preference (C)”)
~~~

By evaluating the policy, one can calculate negative expected free energy for a policy:

~~~
def evaluate_policy(policy, Qs, A, B, C):
 # store expected free energy
 G = 0
 # create copy of our state
 Qs = Qs.copy()
 # loop over policy
 for t in range(len(policy)):
  # get action
  u = int(policy[t])
  # work out expected state
  Qs = B[u].dot(Qs)
  # work out expected observations
  Qo = A.dot(Qs)
  # get entropy
  H = A.entropy()
  # get predicted divergence and uncertainty and novelty
  divergence = F.kl_divergence(Qo, C)
  uncertainty = H.dot(Qs)[0, 0]
  G += (divergence + uncertainty)
 return –G
~~~

And infer an action:

~~~
def infer_action(Qs, A, B, C, n_actions, policy_len):
 # this function generates all possible combinations of policies
 policies = F.generate_policies(n_actions, policy_len)
 n_policies = len(policies)
 # negative expected free energy
 neg_G = np.zeros([n_policies, 1])
 for i, policy in enumerate(policies):
   neg_G[i] = evaluate_policy(policy, Qs, A, B, C)
 # get distribution over policies
 Q_pi = F.softmax(neg_G)
 # probabilites of control states
 Qu = Categorical(dims=n_actions)
 # sum probabilites of controls
 for i, policy in enumerate(policies):
  # control state specified by policy
  u = int(policy[0])
  # add probability of policy
  Qu[u] += Q_pi[i]
 # normalize
 Qu.normalize()
 # sample control
 u = Qu.sample()
 return u
~~~

Putting everything together, we can write a loop to infer beliefs about hidden states:

~~~
# number of time steps
T = 10
# number of actions
n_actions = env.n_actions
# length of policies we consider
policy_len = 4
# set initial state
env.set_initial_state(0)
# reset environment
o = env.reset()
# infer initial state
Qs = F.softmax(A[o, :].log())
# loop over time
for t in range(T):
 # random action
 a = infer_action(Qs, A, B, C, n_actions, policy_len)
 # perform action
 o, r = env.step(a)
 # infer new hidden state
 Qs = F.softmax(A[o,:].log() + B[a].dot(Qs).log())
 # print information
 print(“Time step {}”.format(t))
 env.render()
 plot_beliefs(Qs, “Beliefs (Qs) at time {}”.format(t))
~~~

A straightforward method such as this one could be applied to BCIs to let them use active inference in optimizing their decision-making.

### 1.4. Artificial Curiosity

For artificially intelligent agents to look at their environments and learn from them in a way that a human being would, they develop a kind of artificial curiosity. They can examine and understand which behavior is rewarded based on the outcomes of their actions. Schmidhuber put forward a simple formal theory of fun and intrinsic motivation based on maximizing intrinsic reward for active creation or discovery of novel, surprising patterns[15]. In this sense, artificially curious agents learn to become bored or tired of predictable patterns or behaviors.

Friston’s method of using the free energy principle and predictive coding, the method by which the brain generates and updates a mental model of the environment given sensory input[16][17], doesn’t achieve this, Schmindhuber wrote. By visiting highly predictable states, Friston argued that perception seeks to suppress prediction error by changing predictions and action changes the signals themselves. Instead of learning, Friston’s approach only teaches agents to stabilize and make things predictable. Other methods of using the free energy principle in active inference have included variational Bayes, formal account explaining the relationship between posterior expectations of hidden states, control states and precision[18]. As Friston explained during his session, artificial curiosity falls under a different form of optimality that uses Hamilton’s principle of least action.

## 2. Body

### 2.1. Brain-Computer Interfaces

The BCI is a system that lets a human brain interact directly with an external device for restoring movement [19] or communication[20], for assisting and optimizing tasks[21], and for rehabilitating by enabling the selfregulation of brain activity for therapeutic purposes[22]. The BCI works by collecting information through electroencephalography (EEG) to update an internal model of the user, optimizing interactions via inference about the user, and optimizing interactions by acting upon the user. These behaviors parallel that of an Active Inference system. With Active Inference, the BCI can choose desired outcomes based on the noisy, variable EEG input signal through optimization. By minimizing entropy to anticipate future outcomes, one can use an Active Inference framework to model sequential learning and decision-making in the brain guided by an objective function to minimize free energy when adjusting to signal variability during the signal processing pipeline and modeling and acting in accordance with the causal structure the model determines. The model was shown in the context of a P300-speller, a BCI for communication that uses event-related potentials (ERPs) such as the P300 ERP, an EEG positive deflection around 300 ms after a rare and related event.

By inferring the user’s intentions of which letter to spell (the observation) given a 6 x 6 grid of characters (the action), the machine studies how a user focuses their visual attention and how their brain reacts with a P300. This lets the machine detect a particular ERP in response. The machine flashes the items in repetition until it becomes confident about figuring out which stimulus corresponds to a targeted item[23].

The P300-based BCI has shown improvement with statistical methods have included Bayesian sensor fusions for event detection[24], a classifier with both Fisher’s and Bayesian linear discriminant analysis[25], and a support vector machine classifier[26].

### 2.2. Brain-Computer Interfaces

**Figure 1.**
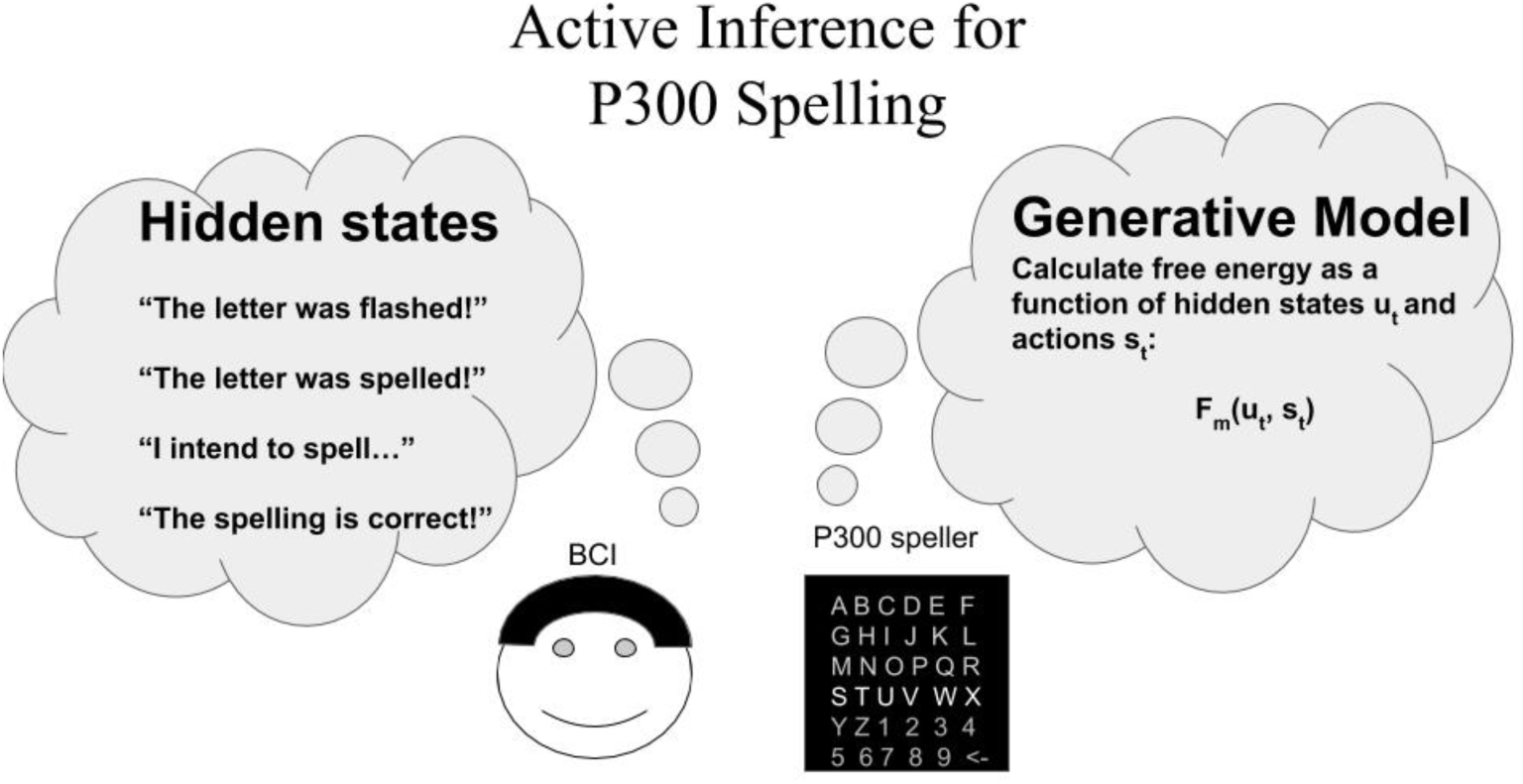
Active Inference for the P300 speller. With free energy as a function of hidden states, the states can be either long term or short term reactions to stimuli. The machine’s actions are based on the generative model of the user. The free energy is, then, a function of hidden states and actions *F*_*m*_(*u*_*t*_, *s*_*t*_).

With this method of Active Inference, one can use a generative Bayesian model of priors that encode predictions over future outcomes with a model m, a joint probability distribution over hidden states *S*, control states (actions) *U*, observations (sensations) *O* and model parameters *S, U*, and *O*. The hidden states are internal representations of the BCI about the etiology of the observations such as the letter a user thinks of that causes the P300 ERP. The Markov model uses the precision parameter *γ* with preferences over future outcomes *C*.

The model works by figuring out which hidden states are most likely to occur by optimizing the expectations with respect to the free energy of the observations. This way, the fully adaptive P300 speller can learn and act optimally in real time by automatically and optimally updating an internal model of the environment. Then, given the speller’s optimized flashing and the corresponding error potentials, the BCI can choose to spell the next probable letter or continue flashing to increase evidence for the target letter. The finite set of hidden states would be: *S* = *s*^(1)^, *s*^(2)^, …, *s*^(*n*)^ with |S| = *n*; in which s maps each trial t onto one element from finite set S such that *s*(*t*) = *s*_*t*_ ∈ *S*, ∀*t* = 1, …, *T*.

If *n* represents the number of possible states (cardinality of *S* at every trial *t*), *T* is the final trial, and *t* is the current one. Only one state out of n possible ones can take place at a time or trial *t*.

By sampling actions from control beliefs and inferring them from observations as well as making the assumption that agents know the realized actions, one can define *U*, a finite set of control states or actions such as which letter to flash next as *U* = *u*^(1)^, *u*^(2)^, …, *u*^(*r*)^, with |*U*| = *r* such that *u* maps *t* onto an element from the finite set *U*. The control states (actions) are then given by *u*(*t*) = *u*_*t*_ ∈ *U* and ∀ *t* = 1, …, *T* in which *r* represents the number of possible states (cardinality of *U* at trial *t*). This lets the agent choose an action out of *r* possible ones.

Similarly, the observations are, then *O* = *o*^(1)^, *o*^(2)^, …, *o*^(*z*)^, with |*O*| = *z* such that *o* maps the trial t onto an element from *O* with *o*(*t*) = *o*_*t*_ ∈ *U*, ∀ *t* = 1, …, *T* for *z*, the number of possible observations (cardinality of *O* at trial *t*).

The Bayesian model is then:

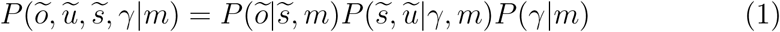

for 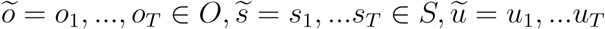. ∀ *U* broken down into the likelihood matrix 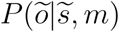, probabilistic transition matrix 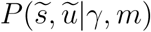, and precision parameter *P*(γ|*m*).

The likelihood matrix:

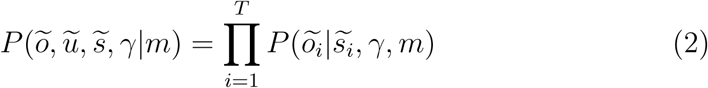

with

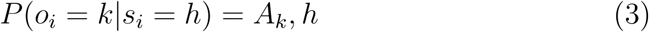

where *A* ∈ ℝ^*zxn*^ so that, for each *h* = 1, …*n* states with a probability to get the *k* = 1, …*z* observation. With the likelihood, the Bayesian model can predict the probabilities of new observations based on prior experience. The probabilistic transition matrix:

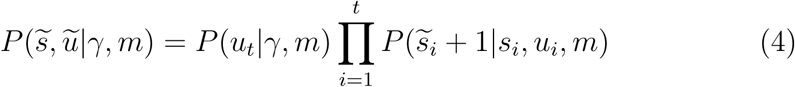

with

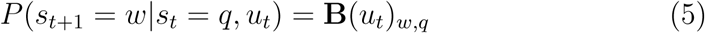

for *w, q* = 1, …*n* and **B**(*u*_*t*_) ∈ ℝ^*nxn*^ for number of hidden states *n*. The putative action, a specific control state that minimizes expected free energy, u_t_ under policy *π* ∈ 1, …*K*. A policy indexes a specific sequence of control states (*ũ*|*π*) = (*u*_*t*_, …, *u*_*τ*_ |*π*) :

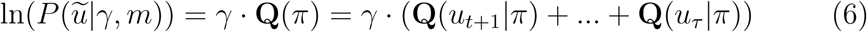

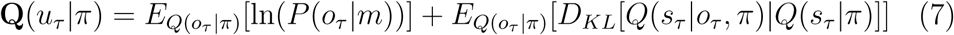

with extrinsic value 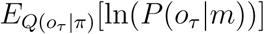 and epistemic value 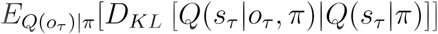, probability distribution *Q*. The precision parameter *γ* is used to minimize expected free energy, *D*_*KL*_ as the Kullback-Leibler (KL) divergence (relative entropy). 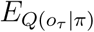 is the expectation of a future outcome, *o*_*tau*_, given policy *π*. The extrinsic value is the maximized preferred final outcome, and the epistemic value is the information maximized achieved by minimizing relative entropy. This way, we maximize information to get to the preferred future outcome.

Using *C*_*τ*_, the prior distribution over the outcomes *P*(*o*_*τ*_ |*m*), we can determine the preference for future outcomes.

The extrinsic value contains *P*(*o*_*τ*_ |*m*) which is the prior distribution over future outcomes, referred to as *C*_*τ*_. So, let *C*_*τ*_ be the preference of future outcomes *o*_*τ*_ ∈ *O*. As part of extrinsic value, it influences the choice of action to reach such desired outcomes. If we consider all available observations from set O as future outcomes then *o*_*τ*_ = *o*^(*z*)^:

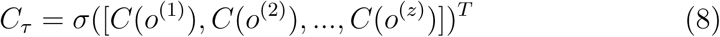

For *σ* softmax (normalized exponential function) of final outcomes that can transform observations to prior probabilities of future outcomes.

The precision parameter is, then:

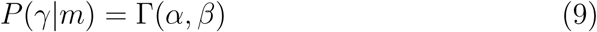

where Γ(*α*) is a gamma distribution of scale parameter a and rate parameter b. If a random variable X follows a gamma distribution then:

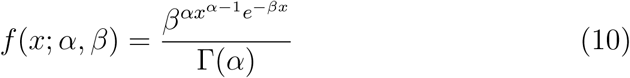

for *x* > 0 and *α*, β> 0.

where Γ (*α*) is the gamma function, as defined:

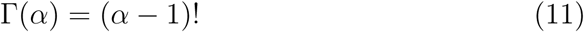

The active inference framework can then be implemented using optimal stopping and flashing while classifying error potential for future outcomes. This way, the machine can act the way the user wants to.

## 3. Conclusion

Active inference provides a way for BCIs to improve their performance in predicting user actions based on how the user interacts with a generative model of the user and their interactions. The general nature of the free energy principle provides a grand, unifying way of understanding how self-organization emerges. The corollary, active inference, then can be used for calculating the epistemic reward, extrinsic value, and other parameters in minimizing free energy and choosing the optimal decision for how the user wants to act. One can describe evidence of specific behavior for active inference in terms of simplicity and accuracy. Paraphrasing Einstein, everything should be made as simple as possible, but not simpler. Minimizing free energy provides a way of achieving this simplicity.

## Declaration of interests

✉ The authors declare that they have no known competing financial interests or personal relationships that could have appeared to influence the work reported in this paper.

□ The authors declare the following financial interests/personal relationships which may be considered as potential competing interests:

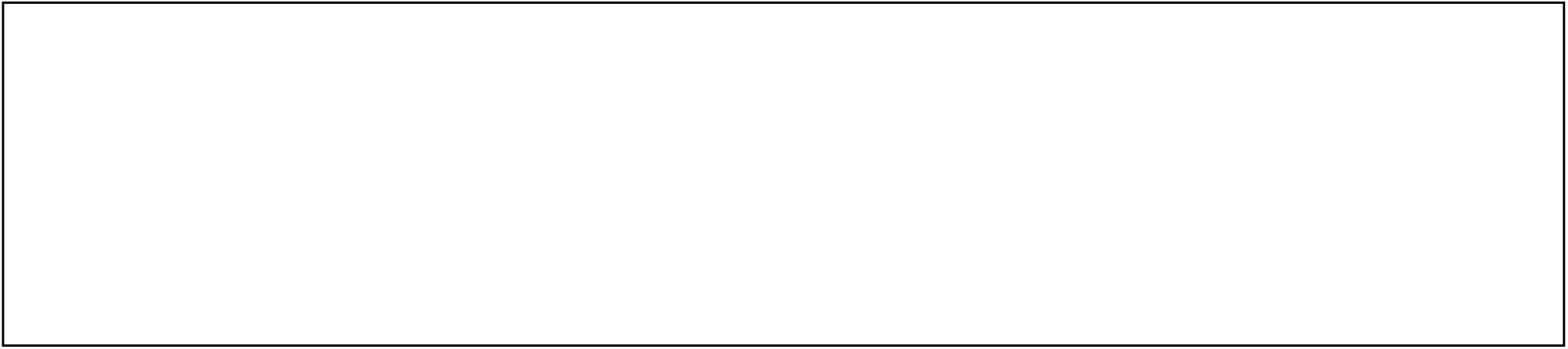

